## Supplementary information for "Mapping tumor spheroid mechanics in dependence of 3D microenvironment stiffness and degradability by Brillouin microscopy"

### Supplementary Figures

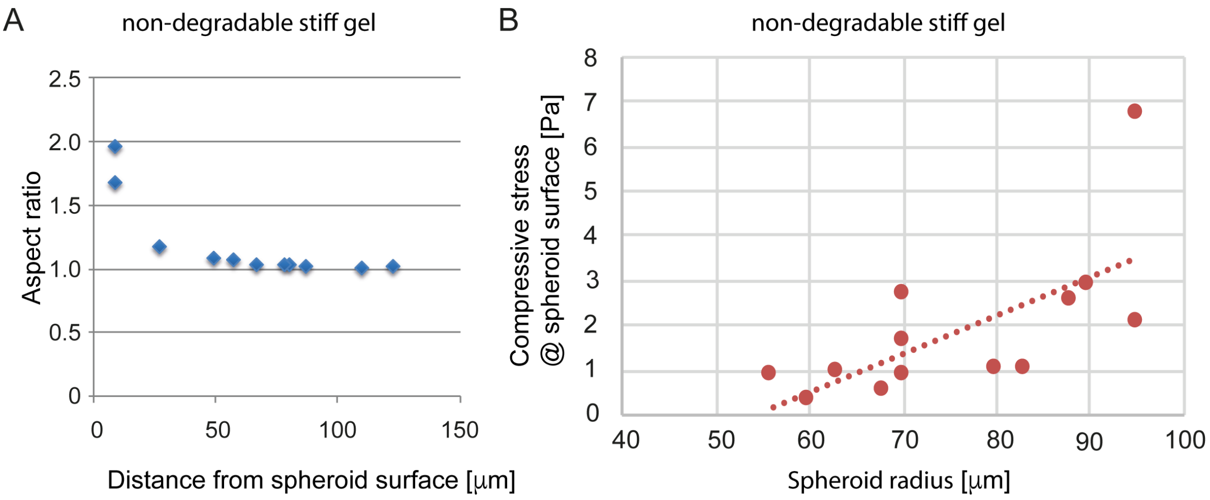


**Supplementary Figure 1. (A)** Example of bead deformations as determined from confocal images using Fiji. Bead aspect ratios were measured in dependence of the distance from the spheroid surface. **(B)** From maximal aspect ratios corresponding radial stresses were calculated. Stresses were plotted over respective spheroid radii (here for stiff non-degradable hydrogels).

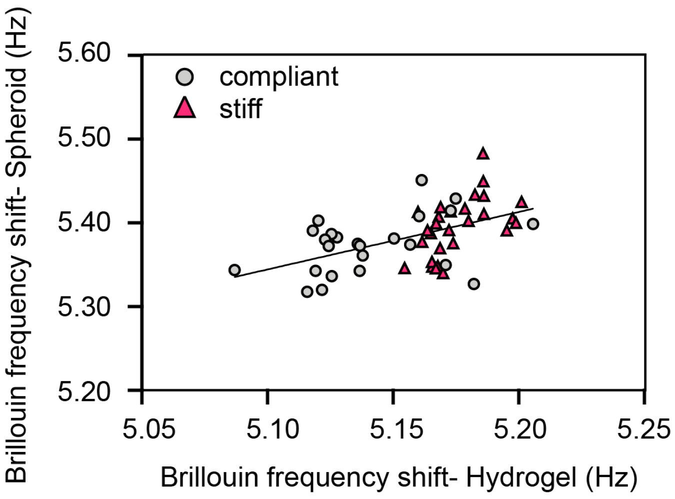


**Supplementary Figure 2.** Brillouin frequency shifts measured for spheroids over corresponding Brillouin frequency shifts obtained from the surrounding hydrogels. A linear trendline is shown.

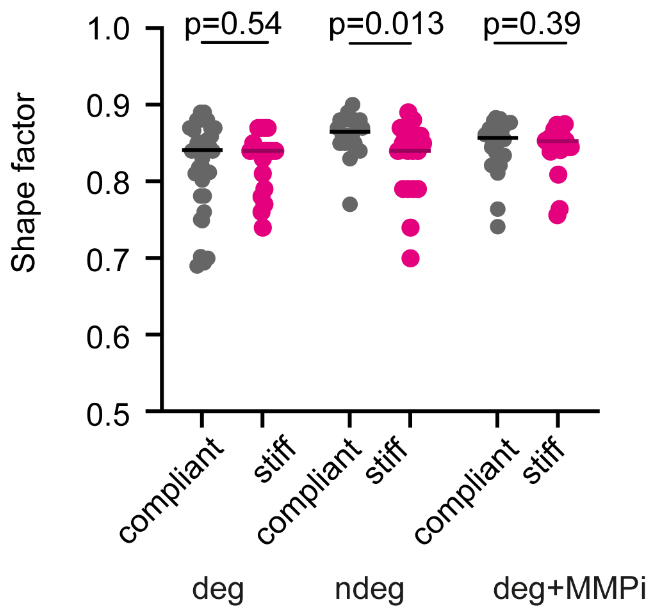


**Supplementary Figure 3. Scatter plots showing shape factors for MCF-7 spheroids formed in compliant and stiff degradable, non-degradable and degradable gels treated with MMPi.** Lines indicate medians. n = 10-20 spheroids each. A Mann-Whitney test was done for statistical analysis. p-values are given.

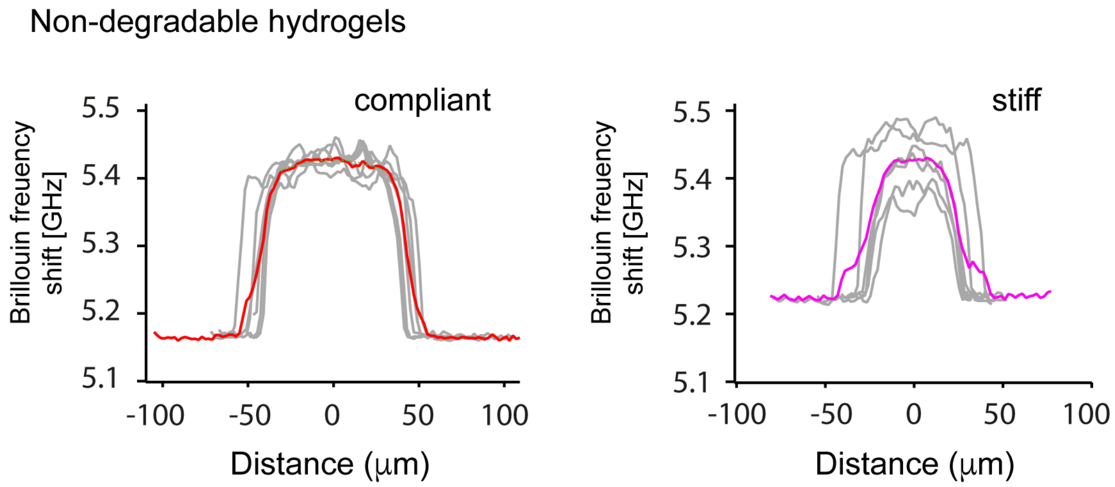


**Supplementary Figure 4.** Averaged line profiles of the Brillouin frequency shift across MCF-7 spheroids grown in compliant and stiff non-degradable hydrogels. Red and pink lines show the calculated averages of grey curves for compliant and stiff hydrogels respectively. Analysis was performed using Igor Pro (Wavemetrics).

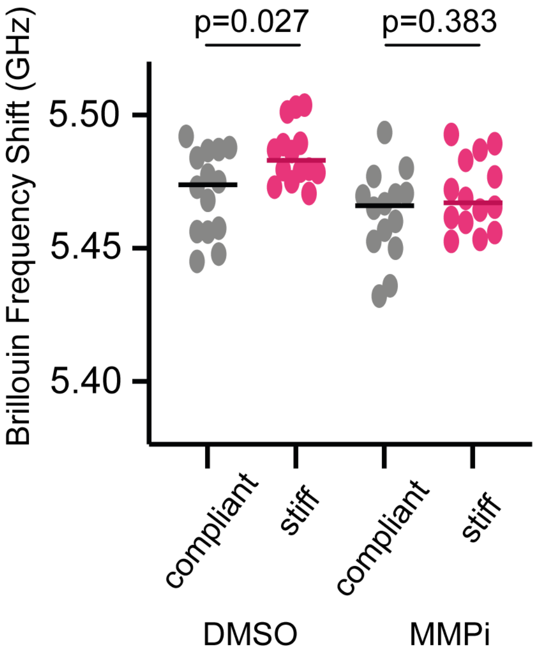


**Supplementary Figure 5. Scatter plots showing Brillouin frequency shifts for MCF-7 spheroids formed in compliant and stiff degradable gels treated with DMSO (vehicle control) and MMPi.** Lines indicate medians. n = 14 spheroids each. A Mann-Whitney test was done for statistical analysis. p-values are given.

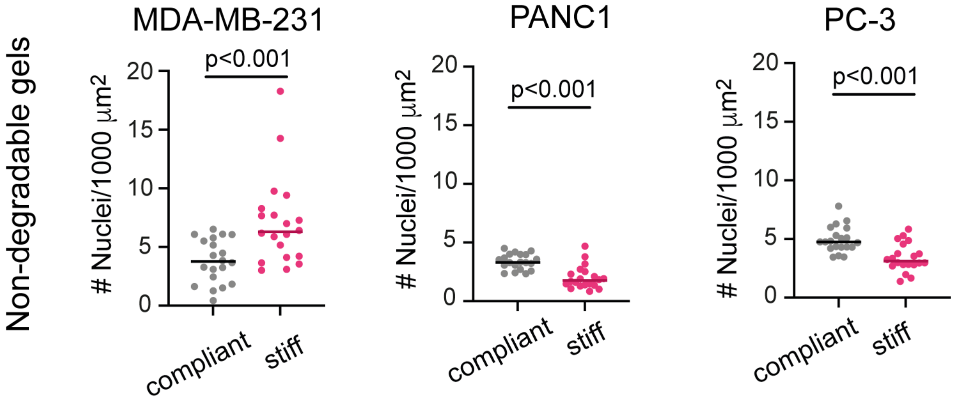


**Supplementary Figure 6.** Scatter plots showing number density of cells per unit area of MDA-MB-231, PANC1 and PC3 spheroids formed in compliant and stiff non-degradable gels as determined using FIJI in confocal microscopy images of Phalloidin/DAPI stained spheroid cultures. Lines indicate medians. n = 20 spheroids each. A Mann-Whitney test was performed for statistical analysis. p-values are given.

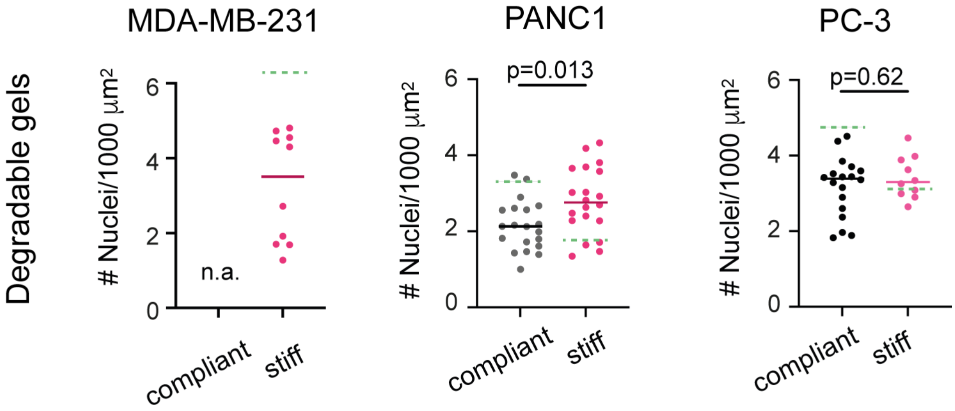


**Supplementary Figure 7.** Scatter plots showing number density of cells/nuclei per unit area of MDA-MB-231, PANC1 and PC3 spheroids formed in compliant and stiff degradable gels as determined using FIJI in confocal microscopy images of Phalloidin/DAPI stained spheroid cultures. Lines indicate medians. n = 10-20 spheroids each. A Mann-Whitney test was done for statistical analysis. p-values are given.

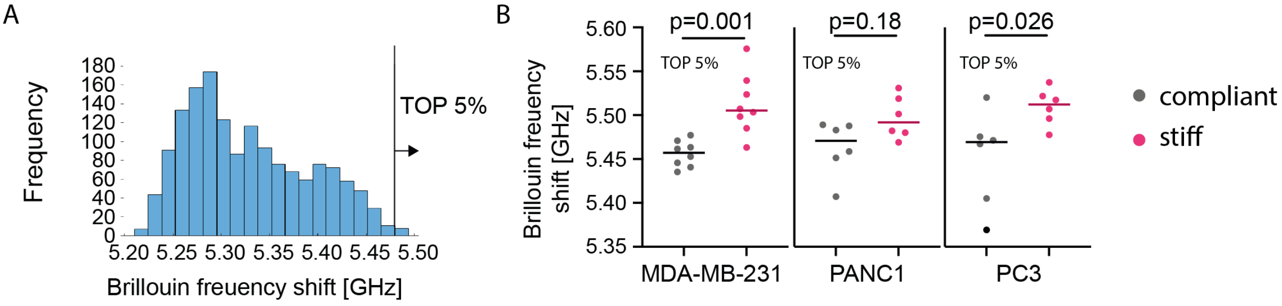


**Supplementary Figure 8.** (**A**) Example histogram showing distribution of Brillouin frequency shift values. (**B**) Scatter plots showing the average of top 5% Brillouin frequency shift values of MDA-MB-231, PANC1 and PC3 spheroids formed in compliant and stiff degradable gels. Lines indicate medians. n = 6-8 spheroids each. A Mann-Whitney test was done for statistical analysis. p-values are given.
